# Commensal gut bacteria convert the immunosuppressant tacrolimus to less potent metabolites

**DOI:** 10.1101/426197

**Authors:** Yukuang Guo, Camila Manoel Crnkovic, Kyoung-Jae Won, Xiaotong Yang, John Richard Lee, Jimmy Orjala, Hyunwoo Lee, Hyunyoung Jeong

## Abstract

Tacrolimus exhibits low and variable drug exposure after oral dosing, but the contributing factors remain unclear. Based on our recent report showing a positive correlation between fecal abundance of *Faecalibacterium prausnitzii* and oral tacrolimus dose in kidney transplant patients, we tested whether *F. prausnitzii* and other gut abundant bacteria are capable of metabolizing tacrolimus. Incubation of *F. prausnitzii* with tacrolimus led to production of two compounds (the major one named M1), which was not observed upon tacrolimus incubation with hepatic microsomes. Isolation, purification, and structure elucidation using mass spectrometry and nuclear magnetic resonance spectroscopy indicated that M1 is a C-9 keto-reduction product of tacrolimus. Pharmacological activity testing using human peripheral blood mononuclear cells demonstrated that M1 is 15-fold less potent than tacrolimus as an immunosuppressant. Screening of 22 gut bacteria species revealed that most *Clostridiales* bacteria are extensive tacrolimus metabolizers. Tacrolimus conversion to M1 was verified in fresh stool samples from two healthy adults. M1 was also detected in the stool samples from kidney transplant recipients who had been taking tacrolimus orally. Together, this study presents gut bacteria metabolism as a previously unrecognized elimination route of tacrolimus, potentially contributing to the low and variable tacrolimus exposure after oral dosing.

## 1. Introduction

Tacrolimus is a commonly used immunosuppressant for kidney transplant recipients as well as patients with glomerular diseases like membranous nephropathy and focal segmental glomerulosclerosis. However, due to its narrow therapeutic index, under-exposure or over-exposure to tacrolimus in kidney transplant recipients increases the risks for graft rejection or drug-related toxicity, respectively.^1^ Maintaining therapeutic blood concentrations of tacrolimus has been difficult in part because tacrolimus pharmacokinetics shows large inter- and intra-individual variability.^2, 3^ For example, tacrolimus oral bioavailability in individual patients ranges from 5 to 93% (average ~25%).^1^ A better understanding of the factors responsible for the variability is crucial for maintaining target concentrations of tacrolimus and improving kidney transplant outcomes.

The human gut is home to over trillions of microbes that can influence multiple aspects of host physiology.^4^ In particular, intestinal bacteria can promote diverse chemical reactions such as hydrolysis and reduction of orally administered drugs, ultimately affecting the efficacy and/or toxicity of drugs.^5-7^ For example, digoxin is converted to the pharmacologically inactive metabolite, dihydrodigoxin, by the gut bacterium *Eggerthella lenta.*^5^ The expression of the enzyme responsible for digoxin metabolism in *E. lenta* is influenced by dietary protein content,^5^ indicating that in addition to the abundance of drug-metabolizing bacteria, diet composition may also govern the extent of drug metabolism in the gut and alter systemic drug exposure. For most clinically used drugs, the involvement of gut bacteria in their metabolism and/or disposition remains unknown.

*Faecalibacterium prausnitzii* is one of the most abundant human gut bacteria (10^8^-10^9^ 16S rRNA gene copies/g mucosal tissue in ileum and colon), taxonomically belonging to the Order *Clostridiales*.^8, 9^ Because of its anti-inflammatory effects, *F. prausnitzii* has been investigated as a potential preventative and/or therapeutic agent for dysbiosis.^10, 11^ We have recently shown that in 19 kidney transplant patients, fecal *F. prausnitzii* abundance positively correlates with oral tacrolimus dosage required to maintain therapeutic blood concentrations, independent of gender and body weight.^12^ It remains unknown, however, whether *F. prausnitzii* is directly involved in tacrolimus elimination in the gut. Herein, we tested a hypothesis that gut bacteria, including *F. prausnitzii*, metabolize tacrolimus into less potent metabolite(s).

## 2. Materials and Methods

### 2.1. Reagents

Tacrolimus was purchased from AdipoGen (San Diego, CA). Casitone and yeast extract were purchased from HIMEDIA (Nashik, MH, IN) and BD (Sparks, MD), respectively. Other components for media were purchased from Thermo Fisher Scientific (Waltham, MA) or Sigma-Aldrich (St. Louis, MO).

Peripheral blood mononuclear cells (PBMC; C-12907) were purchased from PromoCell (Heidelberg, Germany). Phytohemagglutinin (PHA; L1668) and 5-bromo-2′-deoxyuridine (BrdU; B9285) were purchased from Sigma-Aldrich. 3,3′,5,5′-Tetramethylbenzidine (TMB; 34022) was purchased from Thermo Fisher Scientific.

### 2.2. Bacterial strains and growth

*F. prausnitzii* A2-165 was obtained from DSMZ (Deutsche Sammlung von Mikroorganismen und Zellkulturen GmbH). *F. prausnitzii* VPI C13-20-A (ATCC 27766), and *F. prausnitzii* VPI C13-51 (ATCC 27768) were from American Type Culture Collection (ATCC). Other gut bacteria were from Biodefense and Emerging Infections (BEI) Research Resources Repository (Supplemental Table 1). Unless stated otherwise, all the bacterial strains were grown anaerobically (5% H_2_, 5% CO_2_, 90% N_2_) on YCFA agar or broth at 37ºC in an anaerobic chamber (Anaerobe Systems, Morgan Hill, CA), and colonies from the agar plate were inoculated into pre-reduced YCFA broth for preparation of overnight cultures. Optical density at 600 nm (OD_600_) was measured for estimation of bacterial concentration.

### 2.3. Tacrolimus metabolism by gut bacteria

To examine tacrolimus metabolism by gut bacteria, cells of a bacterial strain grown as described above were incubated tacrolimus. Typically, tacrolimus (100 μg/ml) was incubated with bacterial cells in the anaerobic chamber at 37ºC for 24-48 h. Reaction was terminated by adding the same volume of ice-cold acetonitrile. After vortexing for 30 sec, samples were centrifuged at 16,100×*g* for 10 min, and the supernatant was collected for HPLC/UV analysis as described below.

### 2.4. M1 detection

The reaction mixture was analyzed by using HPLC (Waters 2695) coupled with a UV detector (Waters 2487). Typically, 50 μL of a sample was injected and resolved on a C8 column (Eclipse XDB-C8;4.6 x 250 nm; 5 μm) using water (0.02 M KH_2_PO_4_, pH 3.5; solvent A) and acetonitrile (solvent B) as mobile phase with the following gradient: 0-12 min (50% B), 12-17 min (50%-70% B), 17-23 min (70% B), 24-30 min (90% B), and 30-40 min (50% B). Eluates were monitored at 210 nm.

For further verification of M1 production by gut bacteria, the supernatant was also analyzed by HPLC tandem mass spectrometry (HPLC/MS/MS), Agilent 1200 HPLC interfaced with Applied Biosystems Qtrap 3200 using an electrospray ion source. The mobile phase consisted of water with 0.1% formic acid and 0.1% ammonium formate (v/v; solvent A) and methanol (solvent B), and the following gradient was used: 0-2 min (40% B), 2-6 min (95% B), and 6-12 min (40% B). Separation was performed on a Xterra MS C18 (2.1x50mm, 3.5μm; Waters) column at a flow rate of 0.3 ml/min, and M1 was detected at *m/z* 828.5/463.5 in the multiple reaction monitoring mode.

### 2.5. Infrared (IR) and nuclear magnetic resonance (NMR) spectroscopy

IR spectra were acquired on neat samples using a Thermo-Nicolet 6700 with Smart iTRTM accessory. 1D and 2D NMR spectra were obtained on a Bruker AVII 900 MHz spectrometer equipped with a 5 mm TCI cryoprobe. NMR chemical shifts were referenced to residual solvent peaks (CDCl_3_ δ_H_ 7.26 and δ_C_ 77.16). NMR experiments included ^1^H NMR, Distorsionless Enhancement by Polarization Transfer Quaternary (DEPTQ), Homonuclear ^1^H-^1^H Correlation Spectroscopy (COSY), Heteronuclear Single Quantum Coherence Spectroscopy (HSQC), Heteronuclear Multiple Bond Correlation Spectroscopy (HMBC), and ^1^H-^13^C HSQC-Total Correlated Spectroscopy (^1^H-^13^C HSQC-TOCSY).

### 2.6. Kidney transplant recipients’ stool samples

Stool samples were collected from ten kidney transplant recipients during the first month after transplantation at Weill Cornell Medicine and stored at -80°C until analysis. Tacrolimus dosing in each patient was adjusted to achieve a target therapeutic level of 8 to 10 ng/ml. The study protocol for kidney transplant stool sample collection was approved by the Institutional Review Board at Weill Cornell Medicine (protocol number 1207012730).

The microbiota composition of the stool samples was determined using 16S rRNA gene deep sequencing as previously described.^13^ In brief, DNA from stool samples was isolated using a phenol chloroform bead-beater extraction method. The V4-V5 hypervariable region was amplified by PCR and the fragments were sequenced on an Illumina MiSeq (250 x 250 bp). 16S rRNA gene paired-end reads were analyzed using UPARSE^14^ and taxonomic classification was performed using a custom Python script incorporating BLAST^15^ with NCBI RefSeq^16^ as a reference training set.

For the measurement of baseline levels of tacrolimus and M1 in stool samples, an aliquot of stool samples was suspended in PBS (final concentration 20 mg/ml). Also, to measure the capacity of stool samples to produce M1, an aliquot of stool samples was suspended in PBS (10 mg/ml) and incubated with tacrolimus anaerobically for 24 h at 37°C. These samples were mixed with 5 volumes of acetonitrile containing ascomycin as an internal standard. An aliquot (10 µl) was injected into Agilent 1290 UPLC coupled with Applied Biosystems Qtrap 6500. The mobile phase consisted of water with 0.1% formic acid and 10 mM ammonium formate (solvent A) and methanol (solvent B), and the following gradient was used: 0-2 min (20% B), 2-5 min (90% B), and 5-8 min (20% B). Separation was performed on an Xterra MS C18 column (2.1x50 mm, 3.5 µm: Waters) at a flow rate of 0.3 ml/min, with the column temperature set at 50°C. M1, tacrolimus, and ascomycin were detected at *m/z* 828.5/463.4, 821.6/768.6, and 809.5/756.5, respectively, in the multiple reaction monitoring mode. Standard curve (2–100 ng/ml for both tacrolimus and M1) was prepared by spiking tacrolimus and M1 into the stool samples of healthy volunteers.

### 2.7. Statistical analysis

Statistical analyses for comparison between two groups were performed using Student’s *t*-test. A *p-*value ≤ 0.05 was considered statistically significant.

## 3. Results

### 3.1. *F. prausnitzii* potentially metabolizes tacrolimus

To determine whether *F. prausnitzii* is capable of metabolizing tacrolimus, cells of *F. prausnitzii* A2-165 strain were grown to mid-exponential phase in YCFA media, and incubated with tacrolimus (100 µg/ml; 124 µM) anaerobically at 37°C. After 24 h incubation, the mixture was resolved using HPLC and analyzed by a UV detector. The HPLC chromatogram of intact tacrolimus showed multiple peaks, demonstrating tautomer formation as previously reported^17^ (Fig. 1A). For estimation of a concentration of intact tacrolimus, the area of the largest peak at retention time 19.7 min was used. After 24 h incubation with *F. prausnitzii*, the concentration of tacrolimus was decreased by ~50% (Fig. 1B), which was accompanied by appearance of two new peaks (designated M1 and M2, Fig. 1A). The M1 and M2 peaks were not observed when tacrolimus was incubated with boiled *F. prausnitzii* cells (Fig. 1A), indicating that the production of M1 and M2 requires live bacterial cells. Similarly to strain A2-165, two additional strains of *F. prausnitzii* (ATCC 27766 and ATCC 27768) were found to produce M1 and M2 (Supplemental Fig. 1), suggesting that this function is likely conserved in different strains of *F. prausnitzii*.

**Fig 1.**
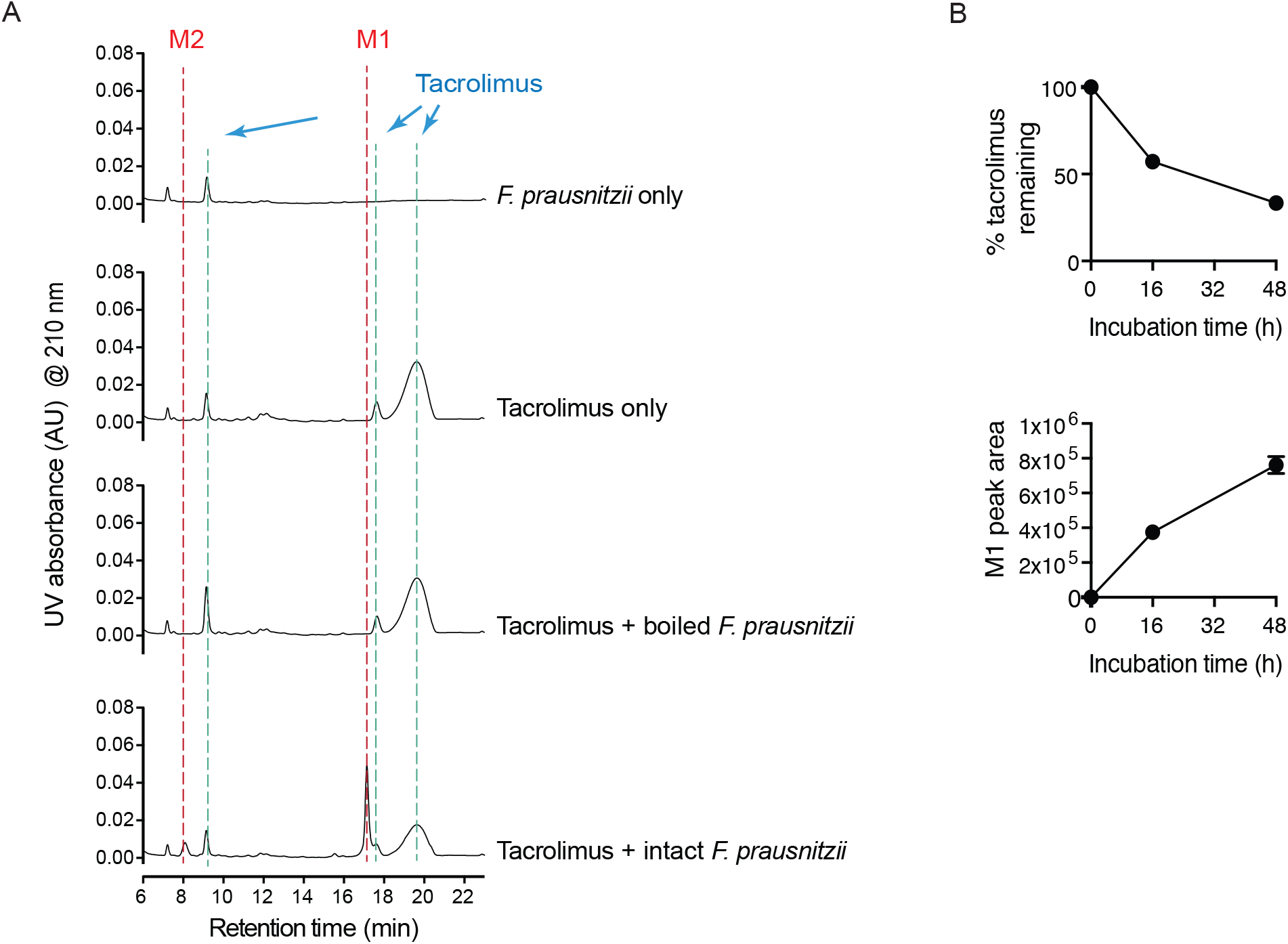
*F. prausnitzii* metabolizes tacrolimus. A, *F. prausnitzii* (OD_600_ 2.6) cultured in YCFA media was incubated with tacrolimus (100 µg/ml) anaerobically at 37°C for 48 h. The mixture was analyzed by using HPLC/UV. B, Time profiles of tacrolimus disappearance and M1 appearance upon anaerobic incubation of tacrolimus (100 µg/ml) with *F. prausnitzii*.

### 3.2. M1 is a C9 keto-reduction metabolite of tacrolimus

To gain insight into the chemical identity of M1 and M2, high resolution mass spectrometry (HRMS) and HPLC/MS/MS experiments were performed. The *m/z* values of M1 and M2 were [M+Na]^+^ 828.4846 and 846.4974, respectively, which are consistent with the formulas C_44_H_71_NO_12_Na (with calculated mass of 828.4874 Da) for M1 (Supplemental Fig. 2) and C_44_H_73_NO_13_Na (with calculated mass of 846.4980 Da) for M2. The calculated formulas suggested M1 to be a reduction product of tacrolimus (i.e., addition of 2H to the parent tacrolimus) and M2 to be a tautomer of M1. The fragmentation pattern of M1 as compared to that of tacrolimus indicated that M1 is likely a keto-reduction product of tacrolimus (Supplemental Fig. 2 and 3).

For structural elucidation, we focused on the major product M1. M1 was mass produced by incubating large amounts of tacrolimus with *F. prausnitzii,* followed by purification using preparative HPLC (Supplemental Methods). The chemical structure of M1 was then determined using various spectroscopic methods. Of note, when the purified M1 was re-injected into HPLC/UV, it resolved into multiple peaks (including one corresponding to M2), indicative of isomerization and/or tautomerization of M1 into M2 (Supplemental Fig. 4). IR spectroscopy further supported that M1 is a product of a carbonyl reduction from tacrolimus (Supplemental Fig. 5). Major differences were observed in the C=O and O-H stretch regions of the IR spectra. NMR spectra showed three major isomers of M1 in CDCl_3_, for which all resonances were assigned (Supplemental Table 3-5). Detailed analysis of 1D and 2D NMR spectra revealed the site of carbonyl reduction at C-9 and the identity of M1 to be 9-hydroxy-tacrolimus (Supplemental Fig. 6-12). In particular, analysis of the DEPTQ spectrum of M1 revealed the absence of the resonances associated with the carbonyl carbon C-9 found in tacrolimus (δ_C_ 196.3 for the major isomer, 192.7 for the minor isomer) (Supplemental Fig. 13). Instead, three resonances consistent with the reduction of the carbonyl at C-9 to an alcohol were observed at δ_C_ 73.0 (isomer I), 68.4 (isomer II), and 69.7 ppm (isomer III). These resonances were associated with protons at δ_H_ 4.02, 4.51, and 4.37 ppm, respectively, in the HSQC spectrum. In turn, the latter resonances showed COSY correlations to exchangeable protons (δ_H_ 4.23, 3.21, and 3.58, respectively). HMBC correlations from H-9 to C-8 and C-10 were observed (Supplemental Tables 3-5), supporting the assignment of M1 as 9-hydroxy-tacrolimus. These results establish the structure of M1 as the C-9 keto-reduction product of tacrolimus (Fig. 2).

**Fig 2.**
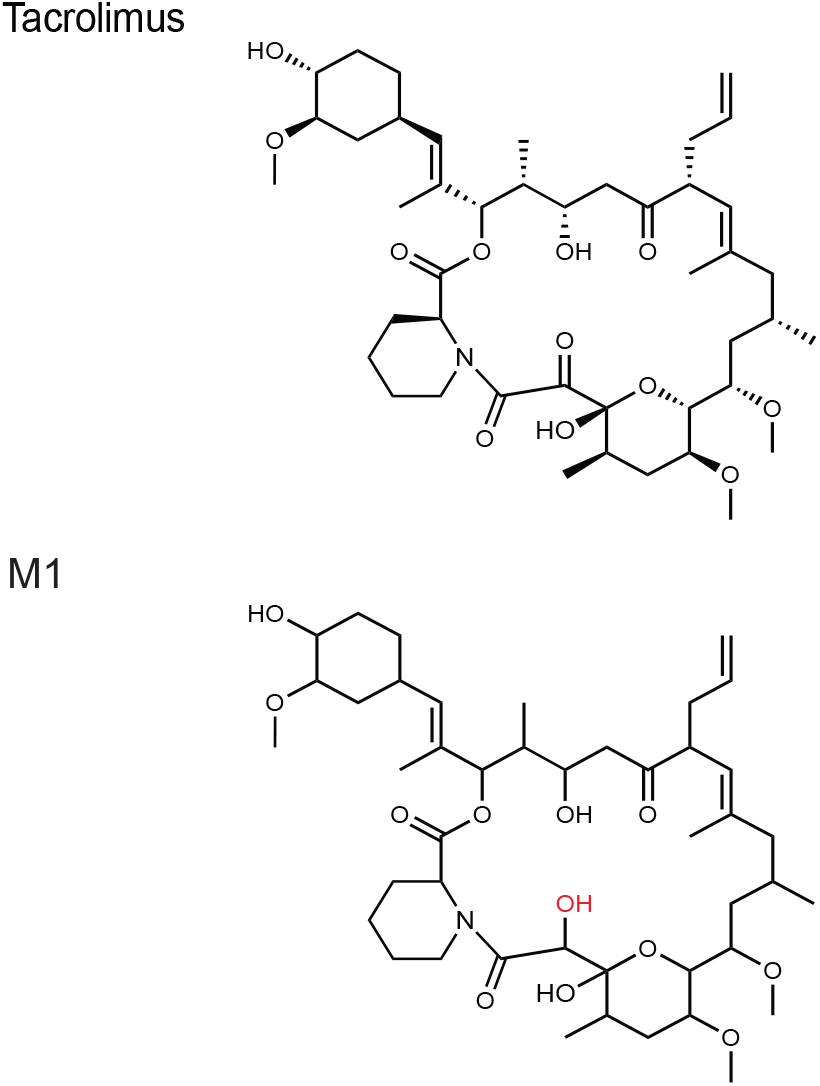
Chemical structures of tacrolimus and *F. prausnitzii-*derived metabolite M1. M1 structure was identified using mass spectrometry and nuclear magnetic resonance spectroscopy.

### 3.3. M1 is a less potent immunosuppressant than tacrolimus

We compared the activities of M1 and tacrolimus by measuring PBMC proliferation after treatment with T-lymphocyte mitogen PHA.^18^ The 50% inhibitory concentration (IC_50_) of M1 was 1.97 nM whereas IC_50_ of tacrolimus was 0.13 nM, demonstrating that M1 was ~15-fold less potent than the parent tacrolimus in inhibiting T-lymphocyte proliferation (Fig. 3A). Tacrolimus is known to exhibit antifungal activity via the same mechanism for immunosuppression.^19^ To further examine the pharmacological activity of M1, an antifungal assay was performed. An aliquot of M1 or tacrolimus was placed onto a lawn of the yeast *Malassezia sympodialis*, and their antifungal activities were estimated based on the size of halo formed. M1 was about 10 to 20-fold less potent than tacrolimus in inhibiting the yeast growth (Fig. 3B), consistent with the results obtained from the PBMC proliferation assay. Taken together, these results demonstrate that M1 is less potent as an immunosuppressant and antifungal agent than the parent drug tacrolimus is.

**Fig 3.**
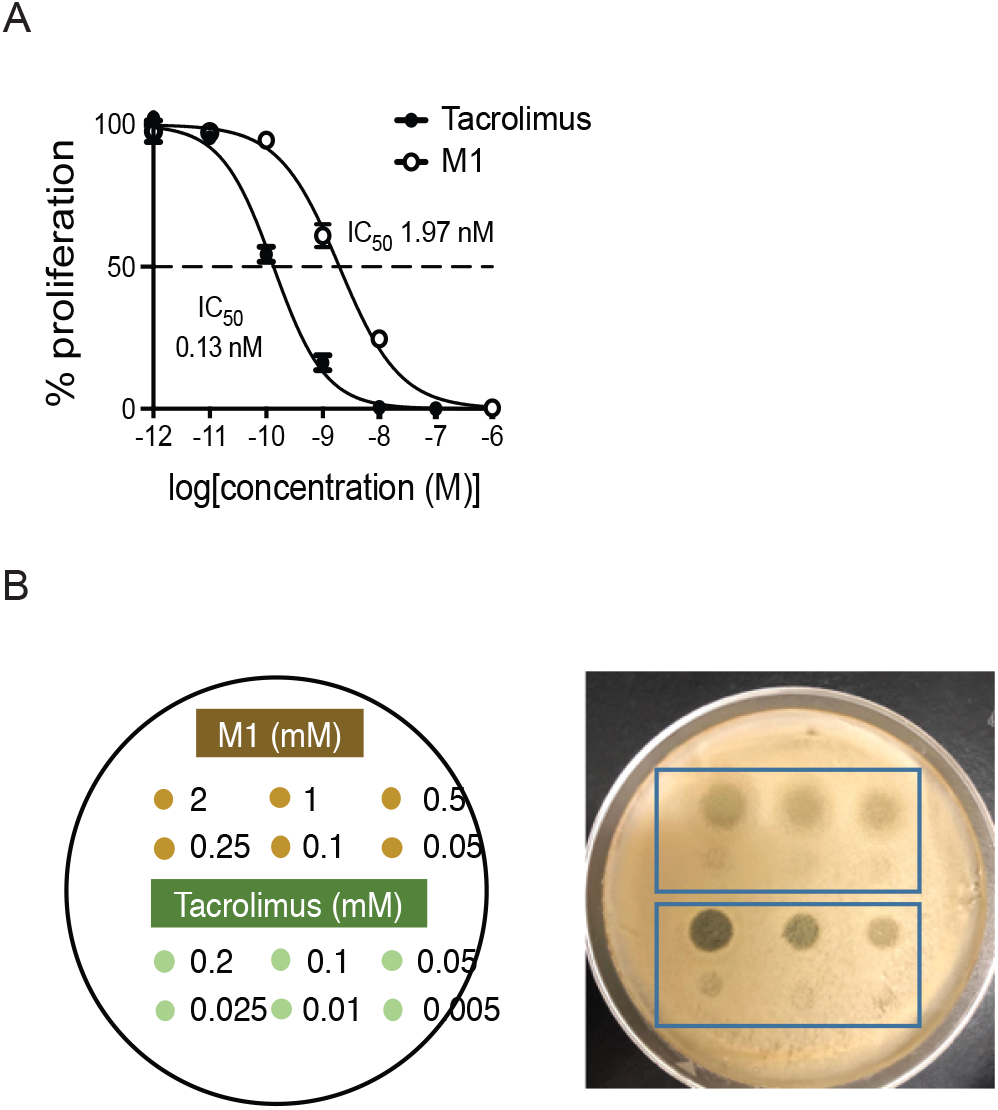
M1 is less potent than tacrolimus as an immunosuppressant and antifungal agent. A, Immunosuppressant activities of tacrolimus and M1 were examined in PBMCs by measuring cell proliferation after treatment with a T-lymphocyte mitogen in the presence of tacrolimus or M1. B, Antifungal activities of tacrolimus and M1 were examined using *Malassezia sympodialis*. The yeast was inoculated on mDixon agar plate. After 1 h incubation, an aliquot of tacrolimus or M1 at different concentrations was placed on the plate as shown on the left panel, and incubated at 37°C for 2 days.

### 3.4. Tacrolimus is metabolized by a wide range of commensal gut bacteria

To determine whether other gut bacteria can produce M1/M2 from tacrolimus, we obtained 22 human gut bacteria from BEI Research Resources Repository (Supplemental Table 1) and tested them for potential tacrolimus metabolism. The tested bacteria include those belonging to major Orders that are known to be highly abundant in the human gut.^8, 9^ Bacteria grown overnight in YCFA media anaerobically were incubated with tacrolimus (100 µg/ml) for 48 h, and the mixtures were analyzed by HPLC/UV. Apparently, gut bacteria in the Orders of *Clostridiales* and *Erysipelotrichales,* but not those in *Bacteroidales* and *Bifidobacteriales* produced M1 (Table 1 and Fig. 4A). To further verify the results, the mixtures were re-analyzed by HPLC/MS/MS which exhibits higher sensitivity than HPLC/UV. M1 production by bacteria in *Clostridiales* was verified (a representative chromatogram of *Clostridium citroniae* shown in Supplemental Fig. 14). M1 production by bacteria in *Bacteroidales* was detectable by HPLC/MS/MS albeit at ~100-fold lower levels than that by bacteria in *Clostridiales* (Supplemental Fig. 14). M1 peak was not detected upon tacrolimus incubation with *Bifidobacterium longum* (Supplemental Fig. 14). The formation of M1 was not observed when tacrolimus was incubated with either human or mouse hepatic microsomes (Fig. 4B; also verified by HPLC/MS/MS, data not shown), suggesting that M1 is uniquely produced by gut bacteria.

**Table 1.**
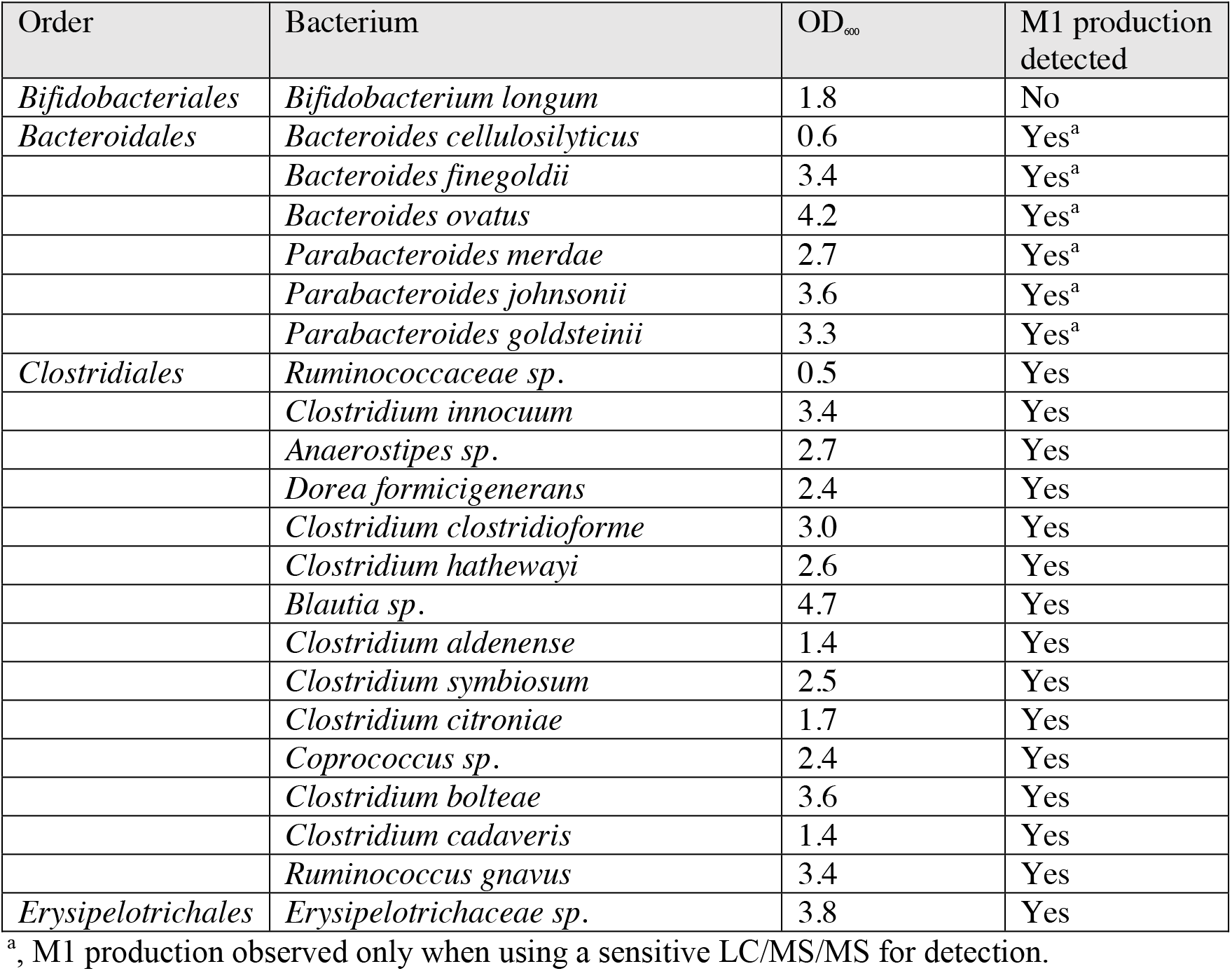
Screening gut bacteria for tacrolimus conversion to M1 in YCFA culture

| Order | Bacterium | OD<sub>600</sub> | M1 production detected |
| --- | --- | --- | --- |
| *Bifidobacteriales* | *Bifidobacterium longum* | 1.8 | No |
| *Bacteroidales* | *Bacteroides cellulosilyticus* | 0.6 | Yes<sup>a</sup> |
|  | *Bacteroides finegoldii* | 3.4 | Yes<sup>a</sup> |
|  | *Bacteroides ovatus* | 4.2 | Yes<sup>a</sup> |
|  | *Parabacteroides merdae* | 2.7 | Yes<sup>a</sup> |
|  | *Parabacteroides johnsonii* | 3.6 | Yes<sup>a</sup> |
|  | *Parabacteroides goldsteinii* | 3.3 | Yes<sup>a</sup> |
| *Clostridiales* | *Ruminococcaceae sp.* | 0.5 | Yes |
|  | *Clostridium innocuum* | 3.4 | Yes |
|  | *Anaerostipes sp.* | 2.7 | Yes |
|  | *Dorea formicigenerans* | 2.4 | Yes |
|  | *Clostridium clostridioforme* | 3.0 | Yes |
|  | *Clostridium hathewayi* | 2.6 | Yes |
|  | *Blautia sp.* | 4.7 | Yes |
|  | *Clostridium aldenense* | 1.4 | Yes |
|  | *Clostridium symbiosum* | 2.5 | Yes |
|  | *Clostridium citroniae* | 1.7 | Yes |
|  | *Coprococcus sp.* | 2.4 | Yes |
|  | *Clostridium bolteae* | 3.6 | Yes |
|  | *Clostridium cadaveris* | 1.4 | Yes |
|  | *Ruminococcus gnavus* | 3.4 | Yes |
| *Erysipelotrichales* | *Erysipelotrichaceae sp.* | 3.8 | Yes |

<sup>a</sup>, M1 production observed only when using a sensitive LC/MS/MS for detection.

**Fig 4.**
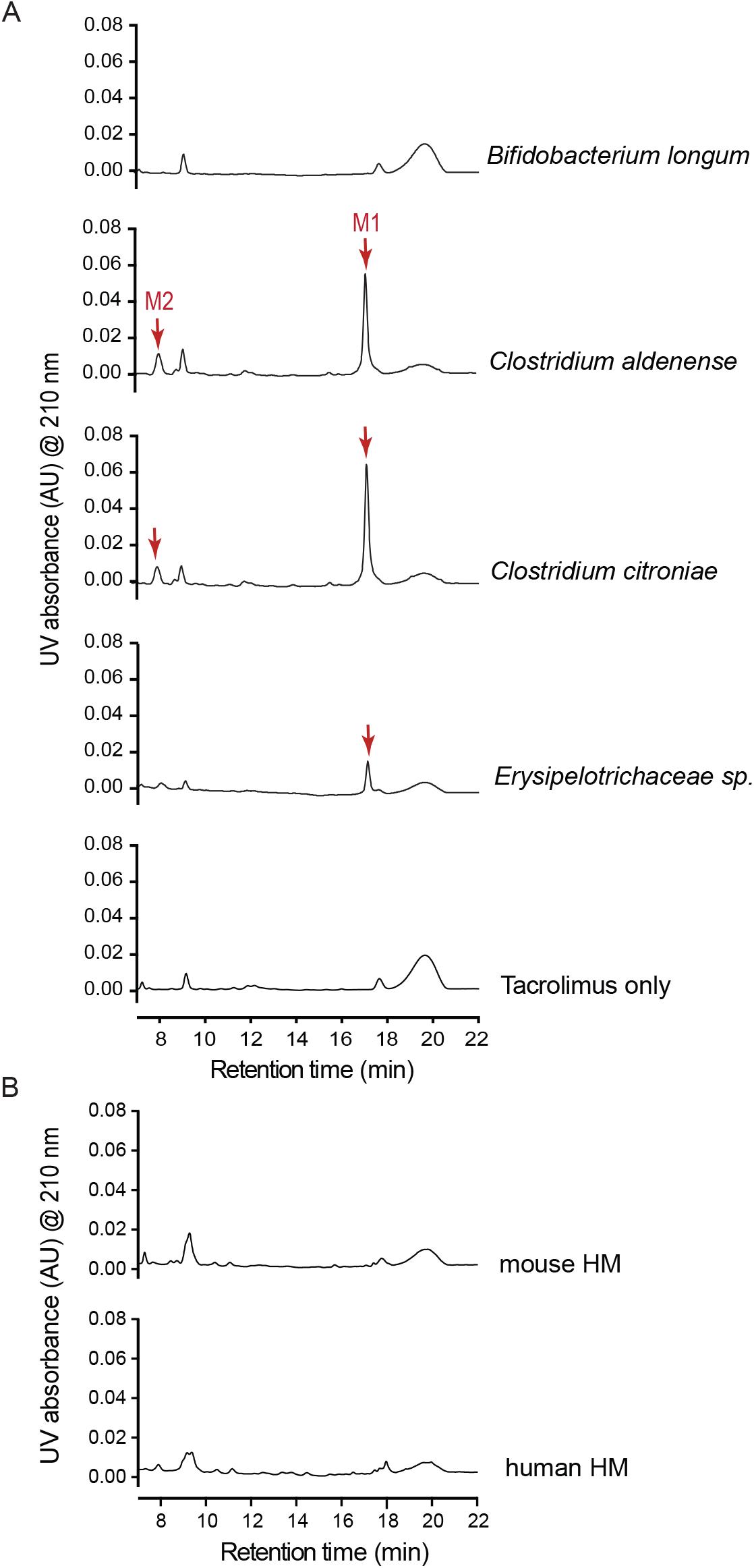
Multiple commensal gut bacteria convert tacrolimus to M1. A, Representative chromatograms of bacteria incubated with tacrolimus. M1 non-producer (*B. longum*) or producers (*C. aldenense, C. citroniae, and Erysipelotrichaceae sp*.) cultured overnight in YCFA media was incubated with tacrolimus (100 µg/ml) anaerobically at 37°C for 48 h. The mixture was analyzed by using HPLC/UV at 210 nm. B, Mouse or human hepatic microsomes (HM; 3 mg microsomal protein/ml) were incubated with tacrolimus (100 µg/ml) at 37°C for 2 h aerobically. The mixture was analyzed by using HPLC/UV.

To examine whether tacrolimus metabolism is indeed mediated by human gut microbiota, fresh stool samples from two healthy adults were incubated with tacrolimus, and M1 production was assessed. Both stool samples produced M1, whereas the control stool samples that were boiled prior to tacrolimus incubation did not (Fig. 5). Taken together, these results show that commensal gut bacteria belonging to different genera metabolize tacrolimus into the less potent M1 metabolite.

**Fig 5.**
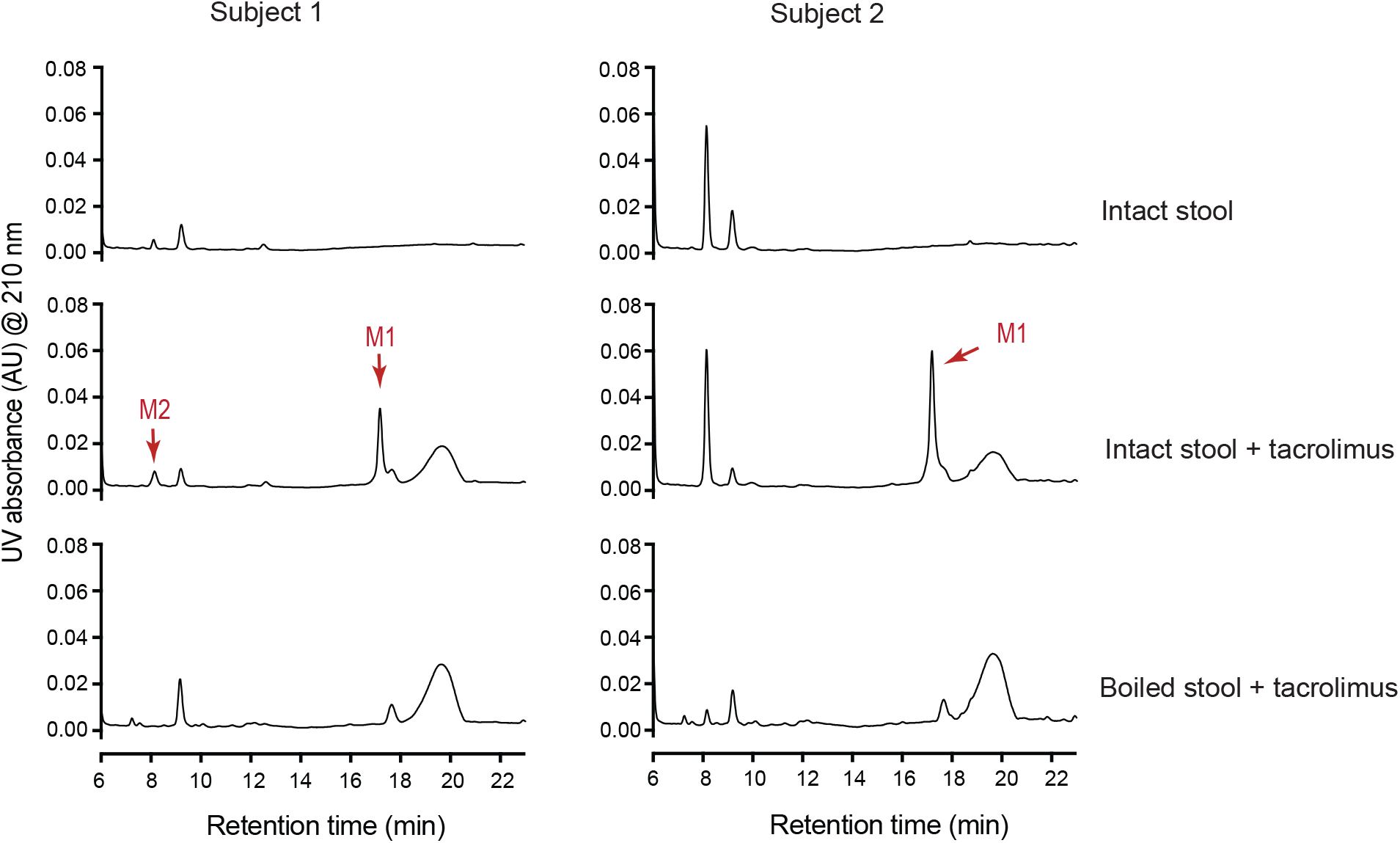
Human gut microbiota convert tacrolimus to M1. Tacrolimus (100 µg/ml) was incubated anaerobically with human stool samples from two different subjects (100 mg wet weight/ml) for 48 h at 37°C. A separate set of sample was boiled for 10 min before incubation with tacrolimus. The incubation mixtures were analyzed by HPLC/UV.

### 3.5. M1 is detected in transplant patients stool samples

*F. prausnitzii* is one of the most abundant human gut bacteria species,^8, 9^ and its fecal abundance was shown to have a positive correlation with oral tacrolimus dosage.^12^ To explore a potential role of *F. prausnitzii* in tacrolimus metabolism in kidney transplant recipients, we evaluated 10 stool samples from kidney transplant recipients who were taking oral tacrolimus (demographic information provided in Table 2 and Supplemental Table 2). Based upon the deep sequencing results of the V4-V5 hypervariable region of the 16S rRNA gene in stool samples, we selected 5 kidney transplant recipients whose stool samples had a relative gut abundance of *F. prausnitzii* greater than 25% (designated as “high *F. prausnitzii”* group) and 5 kidney transplant recipients whose stool samples showed no to little (if any) presence of *F. prausnitzii* (“low *F. prausnitzii*” group). We first determined the baseline levels of tacrolimus and M1 in the stool samples. M1 was detected in three of the five samples from the high *F. prausnitzii* group and one of the five samples from the low *F. prausnitzii* group (Table 2). Because M1 levels were below detection limit for the majority of samples, statistical analysis was not performed. Next, we tested the stool samples of both high and low *F. prausnitzii* groups for the capability of M1 production by incubating each of them with exogenously-added tacrolimus (10 µg/ml) for 24 h. M1 was detected in all 10 samples, but its amount produced was similar between the high and low *F. prausnitzii* groups. The 16S rDNA sequencing analysis of the stool samples revealed that gut bacteria belonging to the Order *Clostridiales* (a main group of bacteria that are expected to produce the majority of M1) were highly abundant in all 10 samples (Table 2). No statistically significant correlation between *Clostridiales* abundance and M1 production was observed (data not shown). Oral tacrolimus dose (to maintain therapeutic blood concentrations) was similar between the two groups (Table 2).

**Table 2.** M1 levels in kidney transplant patients’ stool samples.

| Patient ID | Age (years) | Gender | Post-transplant Day | Tacrolimus oral dose<sup>a</sup> (mg/day) | Fecal abundance of *F. prausnitzii* | Fecal abundance of *Clostridiales* | Baseline tacrolimus level in stool samples (ng/mg stool) | Baseline M1 level in stool samples (ng/mg stool) | M1 production upon tacrolimus incubation (ng/mg stool) |
| --- | --- | --- | --- | --- | --- | --- | --- | --- | --- |
| 1 | 45 | Female | 31 | 9 | 46% | 86% | 0.88 | 0.38 | 5.1 |
| 2 | 56 | Male | 18 | 3 | 39% | 89% | BQL<sup>c</sup> | BQL<sup>d</sup> | 3.5 |
| 3 | 61 | Male | 20 | 5 | 32% | 71% | 0.63 | BQL<sup>d</sup> | 4.5 |
| 4 | 59 | Female | 12 | 6 | 27% | 76% | 0.71 | 0.12 | 2.9 |
| 5 | 50 | Male | 32 | 10 | 26% | 79% | 0.37 | 0.41 | 6.4 |
| 6 | 52 | Female | 28 | 6 | ND<sup>b</sup> | 15% | 0.29 | BQL<sup>d</sup> | 3.5 |
| 7 | 57 | Male | 15 | 3 | ND<sup>b</sup> | 44% | 0.85 | BQL<sup>d</sup> | 4.1 |
| 8 | 71 | Male | 18 | 4 | ND<sup>b</sup> | 95% | BQL<sup>c</sup> | 0.60 | 7.1 |
| 9 | 25 | Male | 27 | 4 | ND<sup>b</sup> | 74% | 0.49 | BQL<sup>d</sup> | 12.6 |
| 10 | 52 | Male | 32 | 6 | ND<sup>b</sup> | 95% | 0.14 | BQL<sup>d</sup> | 11.0 |

<sup>a</sup>, at the time of stool collection
<sup>b</sup>, not detected
<sup>c</sup>, below the quantification limit (i.e., 0.1 ng/mg stool)
<sup>d</sup>, below the quantification limit (i.e., 0.1 ng/mg stool)

## 4. Discussion

In this study, we have demonstrated that a wide range of commensal gut bacteria can metabolize tacrolimus into a novel metabolite M1 (9-hydroxy-tacrolimus). To the best of our knowledge, this represents the first experimental evidence for commensal gut bacteria being involved in the metabolism of tacrolimus.

The extent of M1’s contribution to overall immunosuppression by tacrolimus therapy is unclear. M1 is ~15-fold less potent than tacrolimus in suppressing the proliferation of activated T-lymphocytes and the growth of yeast. This result is consistent with the currently available structure-activity relationships of tacrolimus analogs; modifications at the C-9 position affect the interaction of tacrolimus with its effector protein (i.e., FK506 binding protein 12) and lead to decreased immunosuppressant activities.^20^ While the systemic concentrations of M1 after oral tacrolimus dosing remain to be measured, results from previous tacrolimus disposition studies using a radiolabeled compound^21^ indicate that the blood concentrations of metabolites are likely lower than that of tacrolimus. These results suggest that pharmacological activity originated from circulating M1 is likely less than that from tacrolimus. Of note, certain tacrolimus metabolites (e.g., 13-*O*-demethyltacrolimus), independently of their immunosuppressive activities, cross-react with the antibodies used in the immunoassays for measurement of tacrolimus blood concentrations, leading to overestimation of tacrolimus concentrations.^1, 22^ Interestingly, such overestimation could not be fully explained by the cross-reactivity of currently known tacrolimus metabolites.^22^ Whether the novel metabolite M1 cross-reacts with the antibodies, accounting in part for the overestimation of tacrolimus concentrations, is currently being investigated.

Multiple factors have been reported to contribute to the low and variable bioavailability of orally administered tacrolimus. These include differential expression and/or activity levels of cytochrome P450 enzymes (especially CYP3A4 and CYP3A5 isoforms) and drug transporter P-glycoprotein (P-gp) in the intestine and liver.^1^ Previous pharmacokinetics studies in healthy volunteers and renal transplant recipients have shown that hepatic extraction of tacrolimus is very low (i.e., 4-8%),^23, 24^ suggesting that low oral bioavailability of tacrolimus is mainly due to drug loss in the gut. P-gp-mediated drug efflux and intestinal CYP3A-mediated metabolism were proposed as major contributors to the loss. However, results from drug-drug interaction studies with ketoconazole (a potent inhibitor of CYP3As and P-gp) have shown that oral bioavailability of tacrolimus increases to at most ~30% when co-administered with ketoconazole,^23, 24^ indicating that 70% of oral dose is lost (not reaching systemic circulation) even when intestinal CYP3A and P-gp activities are blocked by ketoconazole. Our results suggest that tacrolimus conversion to M1 in the gut may represent a previously unrecognized pathway of tacrolimus elimination in the gut, and the extent of tacrolimus conversion to M1 in the gut may determine tacrolimus bioavailability. To estimate its overall contribution to systemic tacrolimus exposure, detailed understanding of M1 disposition (e.g., its intestinal absorption and elimination) is warranted.

Our results obtained from testing commensal gut bacteria for tacrolimus metabolism suggest that differences in gut bacterial composition may lead to altered tacrolimus exposure in kidney transplant recipients. Gut bacteria that extensively metabolized tacrolimus into M1 (including *F. prausnitzii*) belong to the *Clostridiales* Order. On the other hand, bacteria in *Bacteroidales* were found to be weak producers of M1 (i.e., detectable only by sensitive HPLC/MS/MS), and *B. longum* in *Bifidobacteriales* did not produce detectable amounts of M1. *F. prausnitzii* is the most abundant gut bacterium at the bacterial species level.^9^ However, we observed no differences in M1 production between high and low *F. prausnitzii* stool samples. Also, we did not observe correlation between *Clostridiales* abundance and M1 production in the stool samples. This may be due to the small number of samples used for this exploratory study and/or the quality of samples non-optimal for enzymatic assays. The presence of multiple factors affecting gut bacterial gene expression *in vivo* such as nutritional status of the gut may further explain why we do not observe a correlation between our *in vitro* culture-based results and *in vivo* abundance of gut bacteria. For example, the amino acid arginine represses the expression of the gene encoding digoxin-metabolizing enzyme in *E. lenta*, thus reducing digoxin elimination by gut bacteria.^5^ Obviously, *in vitro* culture-based systems do not fully reflect the bacterial functions activated in the physiological gut ecosystem. In this regard, our follow-up study is focused on the identification of the bacterial gene(s) responsible for tacrolimus metabolism. Such information will enable us to examine the prevalence and expression of tacrolimus-metabolizing enzymes in gut bacterial community and identify factors such as certain diet or drugs that alter gut bacterial composition and/or gene expression that are specific for tacrolimus metabolism.

In summary, we present the evidence of tacrolimus metabolism by gut bacteria, providing potential explanations for its low oral bioavailability. Tacrolimus metabolism into M1 likely represents a novel elimination pathway that occurs before intestinal absorption of tacrolimus. While the extent of gut metabolism of tacrolimus on variable tacrolimus exposure remains to be determined, our data provide a novel understanding of tacrolimus metabolism and may explain variability in tacrolimus exposures in kidney transplant recipients and patients with glomerular diseases on tacrolimus therapy.

## Author Contributions

All authors designed the study; Y.G., C.M.C., K.-J. W., X.Y. carried out experiments; all authors analyzed the data; all authors made the figures and tables; all authors drafted and revised the paper; all authors approved the final version of the manuscript.

## Acknowledgements and Financial Disclosures

This work was supported by National Institute of Health K23 AI 124464 (J.R.L.) and a grant from the Chicago Biomedical Consortium Catalyst Award C-066 (H.J.).

J.R.L. receives research support from BioFire Diagnostics, LLC. The other authors of this manuscript have no conflicts of interest to disclose as described by the American Journal of Transplantation.

## Supplemental Materials

- Supplemental Methods
- Supplemental Fig 1-14
- Supplemental Table 1-5

## Abbreviations

BEI: Biodefense and Emerging Infections
BrdU: 5-Bromo-2′-deoxyuridine
COSY: Homonuclear ^1^H-^1^H Correlation Spectroscopy
DEPTQ: Distorsionless enhancement by polarization transfer quaternary HMBC Heteronuclear multiple bond correlation spectroscopy
HPLC: High performance liquid chromatography
HRMS: High resolution mass spectrometry
HSQC: Heteronuclear single quantum coherence spectroscopy
IR: Infrared spectroscopy
MS/MS: Tandem mass spectrometry
NMR: Nuclear magnetic resonance spectroscopy
PBMC: Peripheral blood mononuclear cells
P-gp: P-glycoprotein
PHA: Phytohemagglutinin
TMB: 3,3′,5,5′-Tetramethylbenzidine
TOCSY: Total correlated spectroscopy

